# Dynein functions in galectin-3 mediated processes of clathrin-independent endocytosis

**DOI:** 10.1101/2022.07.29.502036

**Authors:** Chaithra Mayya, Hema Naveena, Pankhuri Sinha, Dhiraj Bhatia

## Abstract

Multiple endocytic processes operate in cells in tandem for the uptake of multiple cargoes, metabolites, and signaling molecules that are involved in diverse cellular functions including cell adhesion and migration. The best studied endocytic process involves the formation of a well-defined cytoplasmic coat at sites of uptake made of clathrin and its interacting partners. Galectin-3 (Gal3), an endogenous lectin, binds to glycosylated membrane receptors and glycosphingolipids (GSLs) to drive membrane bending, leading to the formation of tubular membrane invaginations which undergo scission to form a morphologically distinct class of uptake structures, termed clathrin-independent carriers (CLICs). This mechanism has been termed the GlycoLipid-Lectin (GL-Lect) hypothesis. Which components from cytoskeletal machinery are involved in the scission of CLICs remains yet to be explored. In this study, we propose that dynein, a retrograde motor protein, is recruited onto Gal3-induced tubular endocytic pits and provides the pulling force to for friction driven scission. Uptake of Gal3 and its cargoes (CD98/CD147) is significantly dependent on dynein activity, whereas the uptake of transferrin (a marker for clathrin-mediated endocytosis) is only slightly affected upon dynein inhibition. Dynein inhibition also affects cellular organelle distribution, 3D cell invasion and wound healing. Our study thereby reveals functions of dynein in individual and collective cell migration in 2D and 3D that are tightly coupled to endocytic processes in cells.

## Introduction

Endocytosis serves as an important cellular function where cells uptake different components such as nutrients, pathogens, signaling molecules and their receptors (Doherty and McMahon, 2009). Endocytic events can be broadly classified as either clathrin-mediated endocytosis (CME) or clathrin-independent endocytosis (CIE) (Thottacherry *et al*., 2019). Glycosylphosphatidylinositol (GPI)-anchored proteins were among the first identified cargoes that are internalized without the requirement of clathrin or caveolin (Sabharanjak *et al*., 2002). This pathway which is now termed the clathrin-independent carrier (CLIC)-GPI-anchored-protein-enriched early endosomal compartment (GEEC) (CG) pathway(Renard and Boucrot, 2021). The cargoes for CIE include Major Histocompatibility complex (MHC) class I (Dutta and Donaldson, 2015), CD44 (Lakshminarayan *et al*., 2014), Interleukin 2 (IL-2) (Lamaze *et al*., 2001), bacterial Shiga (Renard, Garcia-Castillo, *et al*., 2015), cholera toxins (Day *et al*., 2015), several GPCRs and Receptor tyrosine kinases (RTKs) (Renard, Simunovic, *et al*., 2015; Caldieri *et al*., 2017; Watanabe and Boucrot, 2017; Renard *et al*., 2020).

Galectins have conserved β-galactoside binding sites within the carbohydrate recognition domains (CRDs) which help them recognize the galactose binding partners (Kim *et al*., 2013). Galectins are cytosolic proteins that undergo non-classical secretion. In the extracellular space, they bind to their galactoside containing ligands (Varki *et al*., 2015). Galectins, particularly Galectin-3 (Gal3) can bind to glycosylated cargo (CD98, CD147, integrins…) and glycolipids on the extracellular leaflet of plasma membrane, driving their oligomerization and formation of nanoenvironment for the cellular uptake of glycosylated cargo and glycolipids. This complex of Gal3, receptors and glycolipids induce membrane bending, leading to the formation of tubular membrane invaginations which undergo scission to form endocytic pits without the involvement of clathrin coat proteins. The term ‘glycolipid–lectin (GL-Lect) process has been coined to describe the mechanism of glycolipid- and lectin-dependent endocytosis (Johannes, Wunder and Shafaq-Zadah, 2016).

The cytoskeletal components including Rab GTPases, Rho GTPases and motor proteins, act downstream of the plasma membrane to modulate the dynamics of endocytic processes (Renard and Boucrot, 2021). The microtubule and actin cytoskeletal networks have significant roles in organizing the plasma membrane and in trafficking endocytic carriers (Stephens, 2012). The early endocytic vesicles generated by clathrin mediated endocytosis are transported primarily along actin filaments within the cortical actin network present beneath the plasma membrane (Engqvist-Goldstein and Drubin, 2003). The long-distance movement of cargo and vesicles is mediated by microtubules, to ensure that the cargoes reach the destined intracellular locations (Brown, 1999; Langford, 2002).

Dynein and kinesins transport cargoes along microtubules, whereas the myosin superfamily is actin-based motors. Among the different motors, only dynein and myosin VI (also known as MYO6) mediate retrograde transport from the cell periphery towards the nucleus. Their roles in membrane trafficking are still under investigation. Dynein is shown to elongate Shiga toxin B-subunit (STxB) (Renard, Garcia-Castillo, *et al*., 2015) and cholera toxin B-subunit (CTxB) (Day *et al*., 2015) induced membrane tubules and is also shown to induce friction driven scission of plasma membrane containing BAR domain proteins (Simunovic *et al*., 2017). Here we show that dynein is the key motor protein involved in the uptake and transport of Gal3 mediated endocytic vesicles. Using ATP depletion assay, we show that Dynein provides the pulling force required for the scission of the PM tubules induced by gal3. On inhibition of microtubules and dynein it was seen that gal3 and its cargo uptake was affected thereby shedding new light on the role of retrograde motor protein and cytoskeletal microtubule machinery in the CIE process. We have extended this study to explore the role of dynein inhibition on cellular migration in 2D and 3D systems as well as processes like wound healing, which are acutely regulated by endocytic pathways. Our studies bring out the new functionalities of motor proteins involved in specific cellular endocytic pathways as well as cellular migratory and invasion processes.

## Results

### Dynein inhibition affects galectin-3 and cargo uptake

To study the contribution of dynein in the internalization of Gal3 in CIE processes, we used well known small molecule inhibitors of dynein, Dynapyrazole A (Steinman *et al*., 2017) and Ciliobrevin D (Firestone *et al*., 2012) in mouse embryonic fibroblasts (MEFs). Ciliobrevin D, a pharmacological dynein inhibitor, blocks AAA+ ATPase motor, while dynapyrazole-A blocks the microtubule binding site of dynein without affecting its basal ATPase activity. In the presence of the inhibitors, the uptake of Gal3 was significantly decreased while transferrin, a clathrin endocytic marker gets only partially affected at higher concentrations of this drug (**Figure 1a, b** and supplementary information, **Figure S1 and Figure S2**). This result implies that dynein could be involved in the CIE processes, in particular with the uptake of Gal3, that gets uptaken via CIE. To further establish our hypothesis, dynein was depleted from MEFs using short interfering RNAs (siRNA) against the Dynein heavy chain (DYNC1H1). We found that the Gal3 uptake was indeed affected with endogenous dynein depletion while transferrin uptake was not affected (**Figure 1c,d**).

**Figure 1.**
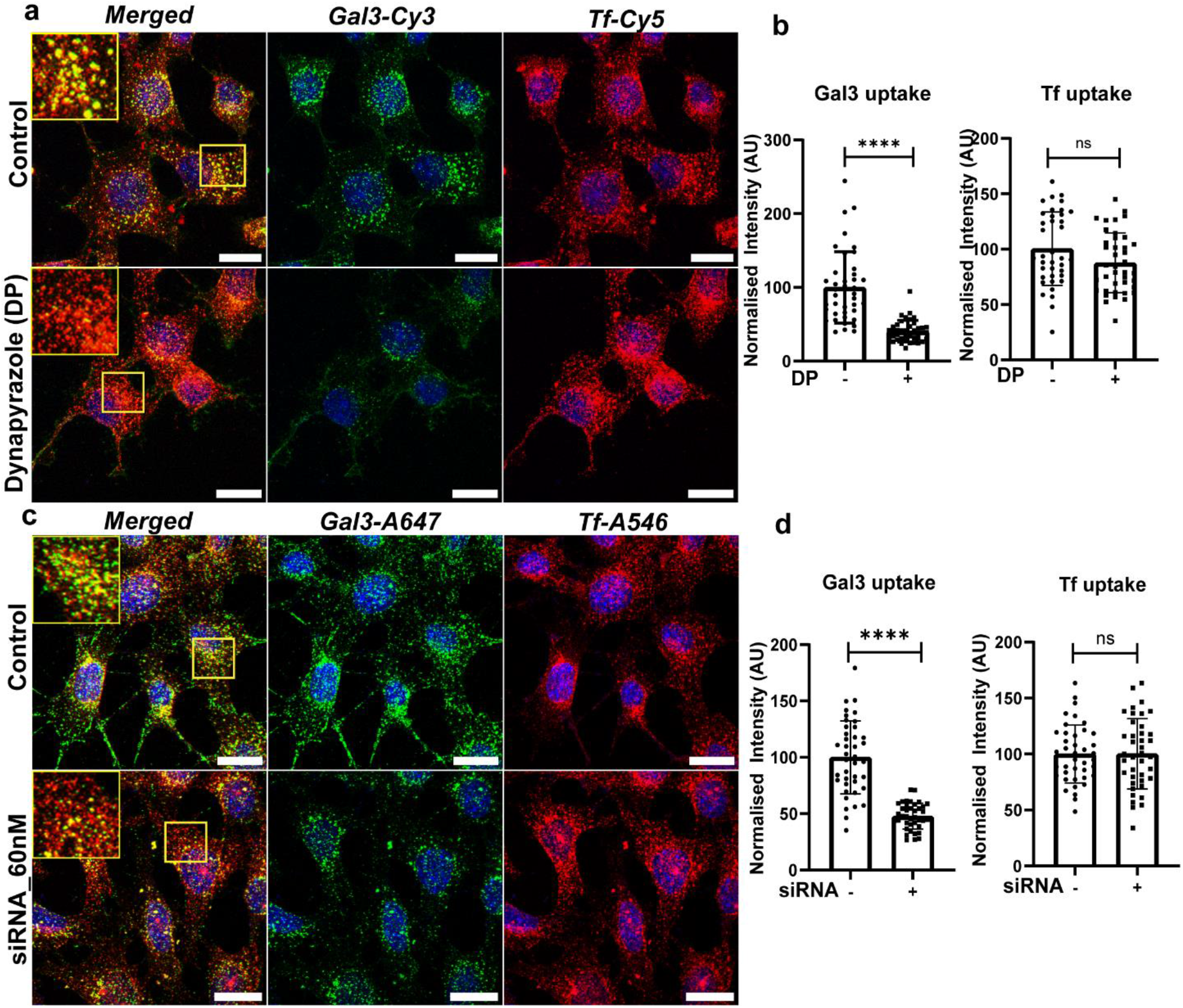
Gal3 uptake is affected upon dynein inhibition. Dynein was inhibited with Dynapyrazole-A/siRNA to check the role of dynein in uptake of Gal3 in MEFs. (a) Dynein was inhibited with 15 μM dynapyrazole for 30 min and Gal3 (Cy3, represented in green) and Tf (Cy5, represented in red) uptake was monitored. Scale bar represents 20μm. (b) Quantification of fluorescence intensity of Gal3 and Tf in control and dynapyrazole treated cells. (c) The dynein was inhibited with siRNA and Gal3 (Alexa 647, represented in green) and Tf (Alexa 546, represented in red) uptake was monitored after 15 mins of uptake. Scale bar represents 20μm. (d) Quantification of fluorescence intensity of Gal3 and Tf in control and siRNA treated cells. Error bars represent the mean **±** S.D. of normalized fluorescence intensity of 35-40 randomly selected cells. Asterisks denote significant difference from the control, with **** p ≤ 0.0001 and ns represents non-significant difference from control.

### Analysis of Gal3 tubulation in plasma membrane invaginations

Using HRP-labelled Gal3 and TEM based imaging, it was earlier shown that Gal3 is detected in tubular plasma membrane invaginations, demonstrating CLIC-like morphology and was microtubule dependent (Lakshminarayan *et al*., 2014). To study the mechanism involved in the bending of the plasma membrane during CIE, we performed ATP depletion-based membrane tubulation assays previously described for STxB, CTxB and Gal3 (Römer *et al*., 2007; Ewers *et al*., 2010; Day *et al*., 2015, Lakshminarayan *et al*., 2014). We found that Gal3 accumulated in the microns-long tubules in the ATP depleted cells and on treatment with CBD these tubule lengths were significantly decreased (**Figure 2a, 2b and 2c**). Dynein motor protein is shown to perform its activity of membrane tubulation even in the decreased ATP conditions (Day *et al*., 2015) and these results indicated that dynein might also be involved in extension process of tubular invaginations induced by Gal3. We observed tubular structures localized at the plasma membrane when immunostained for Dynein heavy chain which reduced upon knockdown of dynein (**Figure 2d, 2e and 2f**).

**Figure 2.**
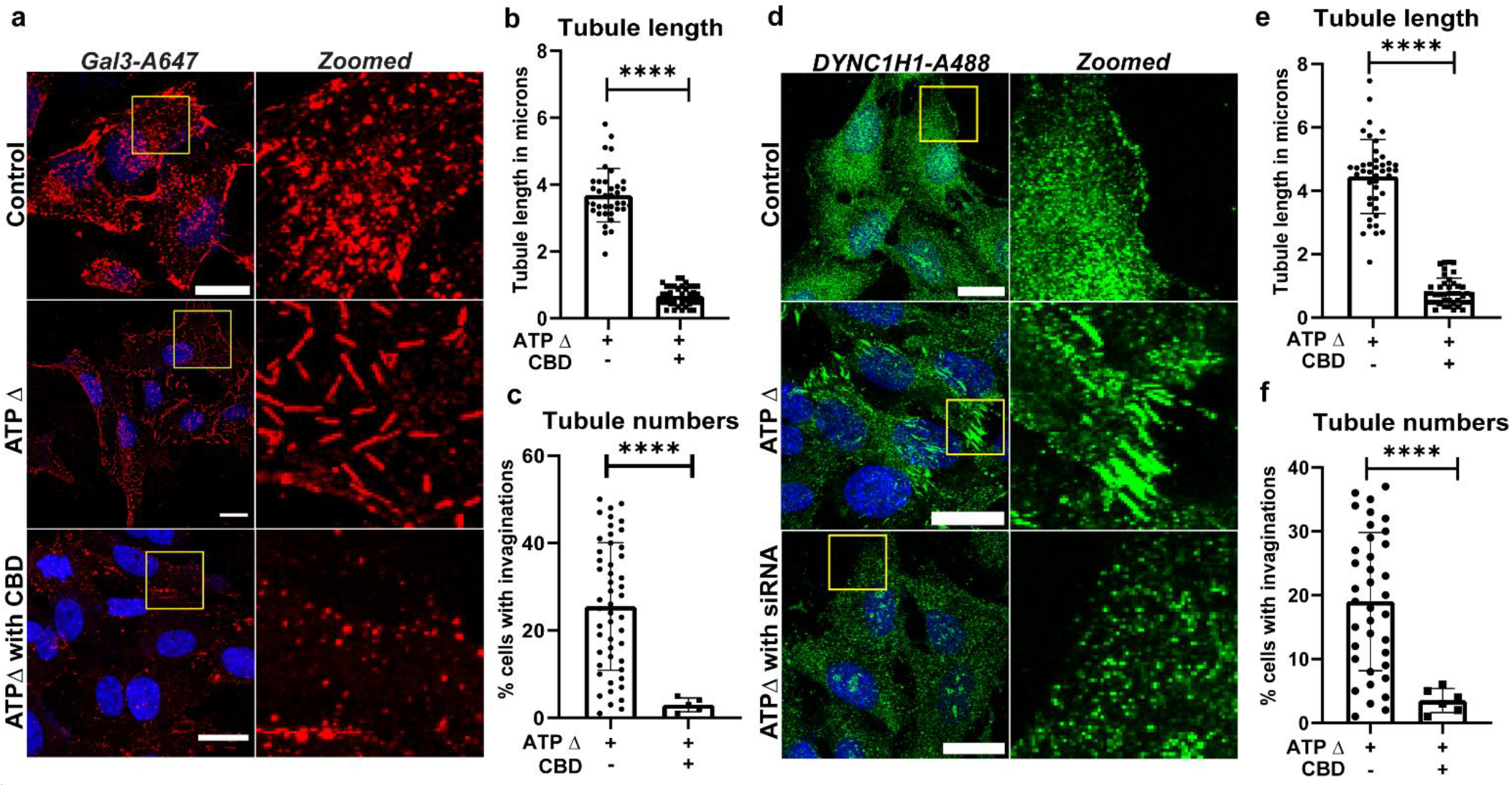
Effect of Gal3 tubulation upon dynein inhibition: Upon ATP depletion there were long tubular structures of Gal3 which were affected on ciliobrevin-D treatment. (a) Long tubular structures of Gal3 (Alexa-647, represented in red) upon ATP depletion for 15 minutes, which reduces on Ciliobrevin-D treatment (15 minutes of inhibitor treatment followed by 15 minutes of ATP depletion). Control cells were not subjected to ATP depletion or CBD (b) Quantification of tubule length for Gal3 in control and Ciliobrevin-D treated cells. (c) The graph represents the number of cells exhibiting optically detectable tubules in control vs CBD treated cells. (d) Confocal images of SUM159 treated with siRNA. In control cells no siRNA treatment or ATP depletion media was added. Long tubular structures are seen in ATP depleted cells stained for DHC (DYNC1H1-A488) while in siRNA treated cells these tubular structures are absent or reduced. The green channel represents DYNC1H1. (e) Quantification of tubule length for DYNC1H1 in ATP depleted and ATP depleted with CBD treated cells. (f) The graph represents the number of cells exhibiting optically detectable tubules in ATP depleted vs CBD treated cells. Error bars represent the mean **±** S.D. of tubular length and number of cells. Error bars represent standard deviation. The tubule length was calculated from 20-25 cells. Student t-test was used for calculating statistical significance with **** p ≤ 0.0001, and ns represents non-significant difference from control. Scale bar is 20μm.

### CD98 and CD147 are Gal3 dependent cargoes

CD98 is a multifunctional protein that plays a role in nutrient uptake and cell-matrix interactions (Devés and Boyd, 2000) while CD147 is a matrix metalloproteinase inducer which associates with monocarboxylate transporter proteins for lactate and pyruvate transportation across the plasma membrane (Iacono *et al*., 2007). It was previously shown that CD98 and CD147 receptors follow the CIE process for their internalization (Eyster *et al*., 2009). It was established that CD44 was Gal3 dependent cargo which got internalized via CIE. We wanted to check if the same was true for other cargoes of CIE such as, CD98 and CD147. We thus used lactose, which is the competitive inhibitor of galectins in general. Lactose binds to the lectin part of galectins, thereby not allowing them to bind to the carbohydrate moiety of cell surface receptors such as CD44, CD98, and CD147, etc. Upon treatment with lactose, there was a drastic decrease in the uptake of CD98 (**Figure 3a,b**) and CD147 (**Figure 3c,d**). We noticed that the distribution of these cargoes in the cytoplasm was affected by Gal3 inhibition using anti-CD98 and anti-CD147 antibodies in SUM159 cells. We observed that the internalization of endogenous CD98 was directly proportional to the increasing levels of Gal3 concentration and were co-localizing at the perinuclear region thereby indicating Gal3 binds to CD98 and stimulates its uptake into the cells (**Figure 3e,f**).

**Figure 3.**
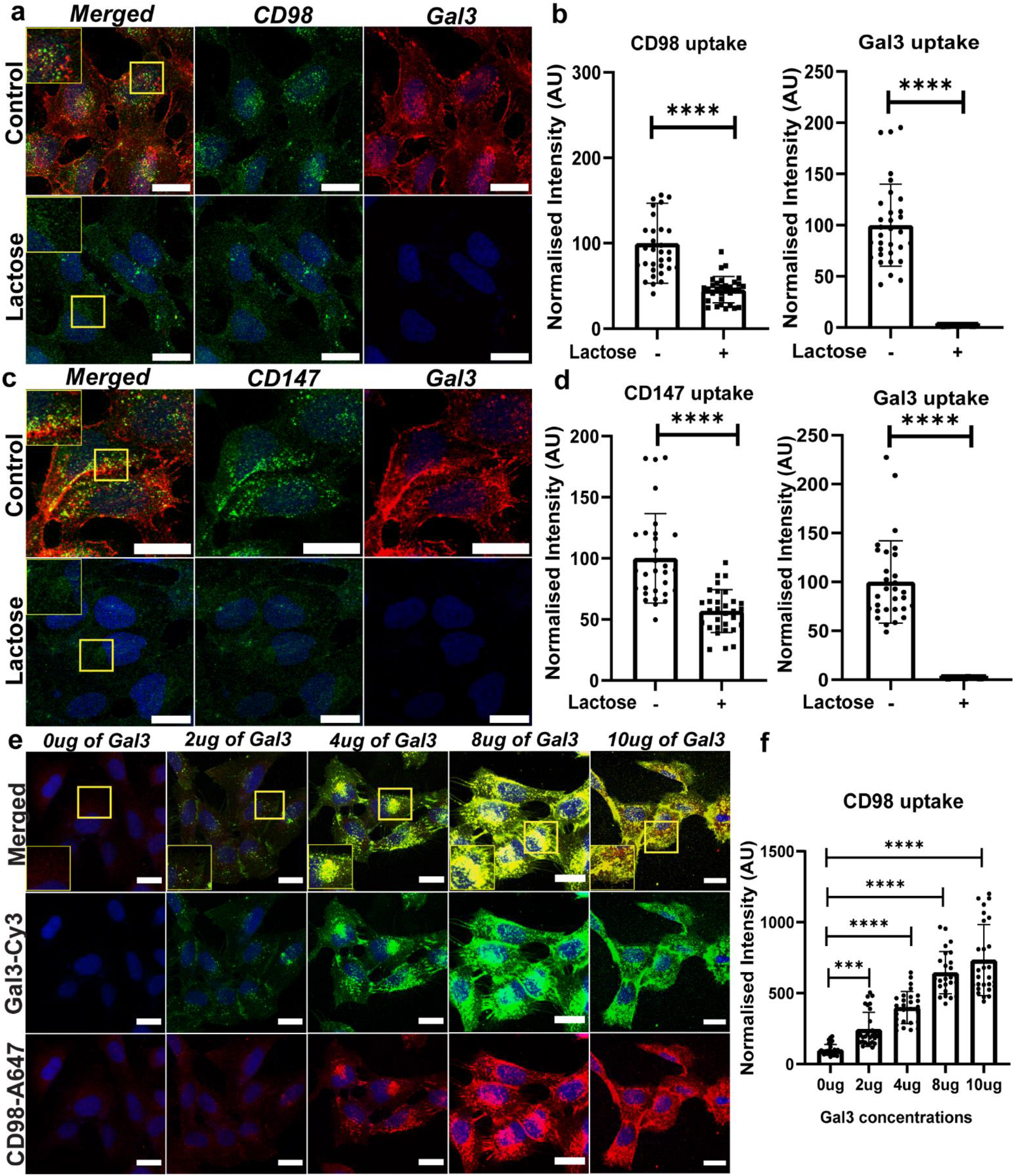
Uptake of membrane receptors like CD98 and CD147 is gal3 dependent: (a) Upon lactose treatment (Gal3 competitor), the internalization of the CD98 receptor is significantly affected similar to Gal3 shown in the figure. (b) Quantification of fluorescence intensity of CD98 in control (without lactose) and lactose treated cells (c) CD147 internalization is also significantly affected on Lactose treatment similar to CD98. (d) Quantification of fluorescence intensity of CD147 in control (without lactose) and lactose treated cells. (Green: Secondary antibody conjugated with Alexa-488; Red: Gal3-A647) (e) Increasing quantities of Gal3 were added to cells and allowed to be uptaken. The internalized fraction of receptor CD98 was immunostained and was quantified as a function of quantity of Gal3 added. Increased perinuclear localization of CD98 upon increased stimulation of Gal3 was observed (upto 10ug). (Green: Gal3-Cy3; Red: Secondary antibody conjugated with Alexa-647). (f) Quantification of increasing fluorescence intensity of CD98 upon increased Gal3 stimulation. Error bars represent the mean **±** S.D. of normalized fluorescence intensity of 40 cells. Asterisks denote significant difference from the control, with *** p ≤ 0.001 and **** p ≤ 0.0001. Scale bar represents 20μm.

### Endocytosis of CD98 and CD147 receptors are also dynein dependent

As established earlier, the above-mentioned receptors are Gal3 dependent following the CIE pathway, we wanted to further check if the uptake of cargos are also dynein dependent similar to Gal3. To test this, we treated the cells with CBD and checked its effect on the uptake of CD98 and CD147. CBD treatment significantly decreased the internalization of CD98 and CD147 similar to that of Gal3 (**Figure S3**). Further to confirm this, we performed the ATP depletion assay to check if these receptors also form tubular structures. In the absence of exogenously added Gal3, the tubules containing CD98/CD147 were very few and small. In presence of exogenously added Gal3, we could observe microns-long membrane tubular invaginations positive for both Gal3 and CD98/CD147; and the tubule size decreased in CBD treatment. This in fact establishes that, CD98 and CD147 are indeed dynein dependent for their internalization (**Figure 4**).

**Fig. 4.**
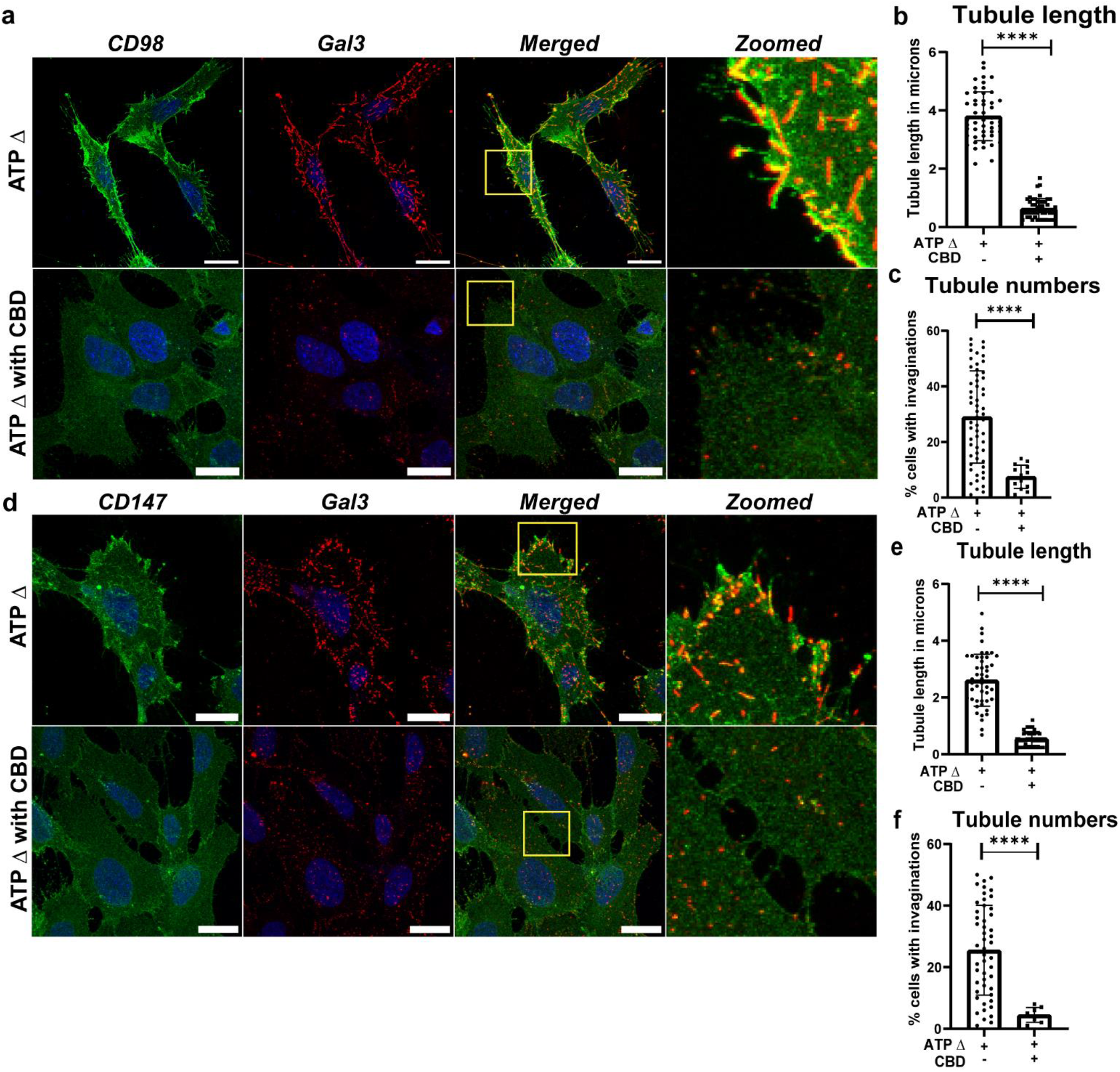
Dynein dependent uptake of CD98 and CD147 in the presence of Gal3: Tubular structures observed for Gal3, CD98 and CD147 under ATP depletion conditions. (a) ATP depletion for 15 minutes in presence of Gal3 resulted in tubulation of CD98 just like Gal3. This tubulation is decreased in terms of size and number of tubules by CBD treatment, indicating dependence of CD98 on dynein. (b) Quantification of tubular length of CD98 in ATP depleted cells (control) versus CBD treated cells. (c) Quantification of number of cells exhibiting CD98 tubulation. (d) ATP depletion in the presence of Gal3 resulted in tubulation of CD147 as well just like Gal3 and CD98, indicating that CD147 is also dynein dependent. (e) Quantification of tubular length for CD147 in ATP depleted and CBD treated cells. (f) Quantification for number of cells showing CD147 tubules. Prior to ATP depletion the cells were treated with the CBD for 15 mins. Scale bar represents 20 μm. Error bars represent the mean **±** S.D. of tubule length and number of tubules. Number of cells taken for counting are 40 cells. Asterisks denote significant difference from the control, with **** p ≤ 0.0001.

### Endocytosis of other galectins also requires dynein activity

There are different types of galectins with different specificities and binding affinities for membrane receptors. We wanted to check if the dynein dependency of endocytic vesicles is only redundant to gal3 induced vesicles or does it apply to other galectins as well. Galectin 8 (Gal8) is another one of the key players in providing the extracellular membrane reorganization inducing membrane bending and formation of pits in case of CD166 uptake, assisted by glycosphingolipids and Endophilin A3 (Renard *et al*., 2020). Like Gal3, Gal8 is also inhibited by lactose (**Figure S4**). We found that inhibition of dynein also affected Gal8 uptake in cells (**Figure 5**). This indicated that dynein probably has a bigger role to play in CIE involving other lectins as well. Similarly, the activity of dynein in terms of providing the pulling force in the other similar membrane tubules induced by bacterial toxins like Cholera toxin B-subunit (CTxB) has been established, indicating the role of dynein in most of the clathrin independent pathways.

**Figure 5.**
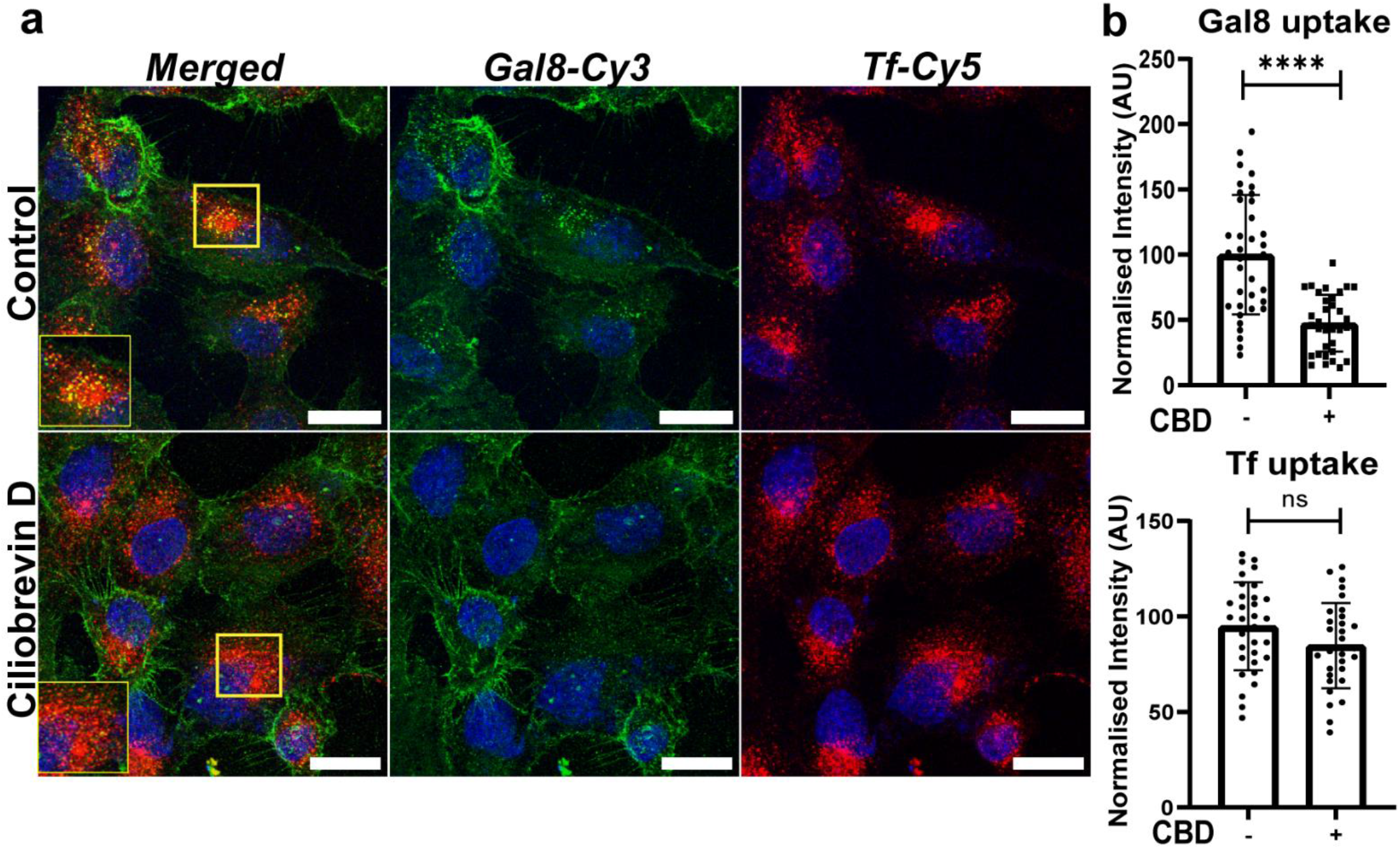
Role of Dynein in Gal8 uptake. (a) Upon CBD treatment, a significant decrease in Gal8 is observed while Tf remains unaffected. Scale bar represents 20μm. (Green: Gal3-Cy3; Red: Tf-A647). (b) Quantification of fluorescence intensity of Gal8 and Tf in control vs CBD treated cells. Error bars represent the mean **±** S.D. of normalized fluorescence intensity of 35-40 cells. Asterisks denote significant difference from the control vs CBD treated, with and **** p ≤ 0.0001 and ns represents non-significant difference from control.

### Dynein inhibition affects 2D wound healing and 3D cell invasion processes

Endocytic trafficking is closely linked to cell migration and invasion (Maritzen, Schachtner and Legler, 2015; Paul, Jacquemet and Caswell, 2015). Clathrin independent endocytic processes are involved in cell migration in migratory cells (Howes *et al*., 2010; Shafaq-Zadah, Dransart and Johannes, 2020). Dynein is a multifunctional motor having multiple roles in regulating the organelle positioning (Burkhardt *et al*., 1997; Lam *et al*., 2010). However, its role in cell migration and invasion is still unexplored. Hence, we wanted to check if apart from playing the role in regulating endocytic pathways, if Dynein played any role in cellular pathways like cell migration and invasion.

We extended our study to check physiological effects of dynein inhibition in 2D and 3D cell culture systems. We used 2D scratch assay to create a wound in migratory Rpe1 (retinal pigmented epithelial) cells to represent a 2D system with or without dynein inhibition. As compared to the control, we saw less would closure in CBD treated cells (**Figure 6a,b**). This indicated that dynein would be playing a role in the migratory aspect of the cells. For 3D invasion studies, we used 3D spheroids as a model system from MDA-MB-231 cells. The invasion index of CBD treated spheroids was significantly reduced as compared to the control (**Figure 6c,d**). In both wound healing and 3D cell invasion, we used cells treated with exogenous Gal3 as positive control; and found that Gal3 enhances the both the wound closure as well as invasion of the cells in 3D, indicating that GL-LECT process might be activating the cell migration and invasion via Gal3 mediated endocytosis of the membrane cargos (**Figure 6a-d**). These above results suggest that dynein is playing a role 2D wound healing as well as 3D invasion processes, which indeed shows the physiological relevance of this motor protein.

**Figure 6.**
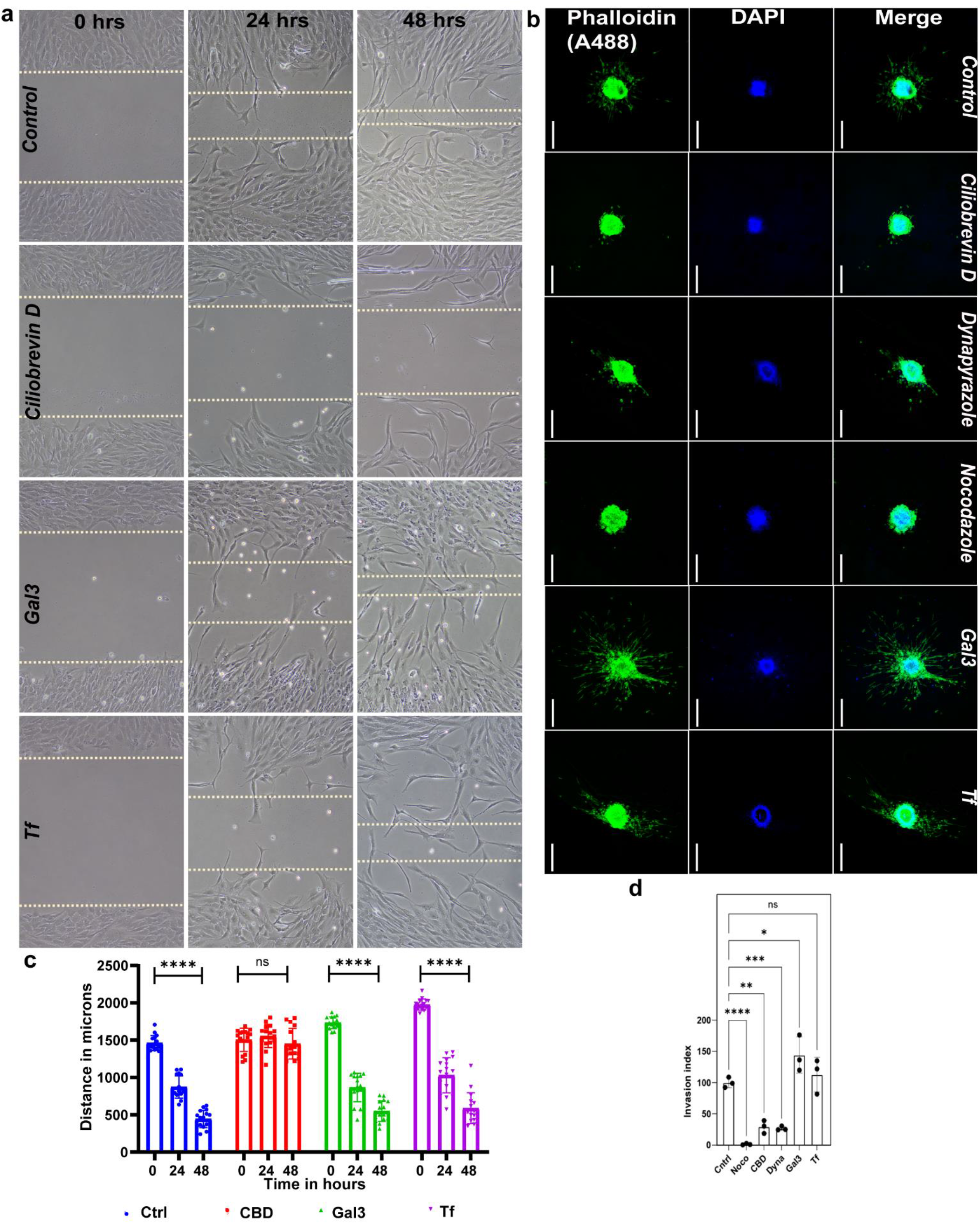
Effect of dynein inhibition on Wound healing and invasion: (a) Scratch assay performed to determine the effect of dynein inhibition on wound healing. The gap size is still retained in CBD treated cells (RPE1 cells). The wound closure is evident in Gal3 treated cells. (b) Invasion assay in 3D spheroids using MDA-MB-231 cells. The CBD and dynapyrazole treated cells have less invasion index compared to control and gal3 treated spheroids. (c) Distance of the wound closure is calculated from 16 different regions per condition and quantified. Quantification of wound size created due to scratch made. ****p≤ 0.0001 shows significant difference between control and treated cells. 2-way anova was used to derive statistical significance. (d) Statistical analysis of the invasion index in 3D spheroid model with ****p≤ 0.0001, ***p≤0.001, **p≤0.01 and *p≤0.05 significance. Student t-test was used for comparing between control and each set of treated spheroids.

## Discussion and conclusions

Our studies show that dynein provides an important mechanical pulling force for the uptake of Gal3 mediated tubular scission during clathrin independent endocytosis. Recent evidence shows that dynein and microtubules are involved in the uptake of CTxB (Day *et al*., 2015) and STxB (Renard, Simunovic, *et al*., 2015). Thus, it’s getting established that dynein and microtubule-dependent membrane tubular structures are a general characteristic of clathrin-independent carriers. We see similar tubular structures for Gal3 as seen for CTxB under limited availability of ATP (**Supplemental information, figure S5**). One other retrograde motor kinesin-14 could be involved in these processes, being a retrograde motor protein utilizing microtubules network. Dynein provides an internal pulling force beneath the plasma membrane which could be sensed by curvature sensing proteins such as Endophilins to promote scission. There was a considerable decrease in Gal3 uptake in Myosin VI inhibited cells which could be stating the involvement of other retrograde motor protein although this has to be further explored such as knockdown studies.

The exact mechanism of how these motor proteins is recruited to the plasma membrane invagination and their regulation in CIE is still unclear. There could be specific adaptor proteins and regulators of dynein playing a pivotal role such as Hook proteins, Rab GTPases, Arfs and so on. The role of Hook1, a known dynein adaptor and microtubule tethering protein is involved in CIE processes for cargoes such as CD44, CD98 and CD147 (Maldonado-Báez *et al*., 2013; Higashi *et al*., 2022). However, we could not find the co-localization of Hook1 and Gal3 tubules in ATP depleted cells (**supplementary figure S6**), yet this aspect of study is still under exploration. There could be other regulatory proteins of dynein which could be involved in Gal3 uptake as well as other galectins.

Our study also shows the role of the actin cytoskeleton in Gal3 uptake. Perturbation of actin showed significant reduction in Gal3 uptake which could be highlighting the importance of cortical actin which could be playing a role beneath the plasma membrane as well as at the sites of plasma membrane invaginations. This could be due to alteration in actin dynamics could decrease the plasma membrane tension which could be favoring the tubulation process in CIE process as hypothesized in previous results (Day et al., 2015). On the basis of CTxB results, we also conclude that not all CLICs are Gal3 dependent. There could be other galectins involved in the formation of CLICs that would be cargo dependent. The plasma membrane components such as glycosphingolipids would also be contributing factors (Lakshminarayan *et al*., 2014). However, we did find that CD98 and CD147 are Gal3 dependent cargoes and they also require dynein for their endocytosis. However, we still need to explore the other contributing factors for their uptake such as role of actin and microtubule, role of GSLs. Also, it would be interesting to explore how Gal3 interacts with these glycosylated cargoes that leads to the uptake of these receptors in dynein dependent manner.

In conclusion, we propose that dynein motor protein pulls the plasma membrane invagination induced by GL-LECT. Cortical actin and its associated factors such as myosin VI would also be modulating the cell surface thereby assisting in the tubulation of plasma membrane. There could be synergistic effects of microtubule and actin dynamic operating beneath the plasma membrane which affects CIE processes (**Figure 7**).

**Figure 7.**
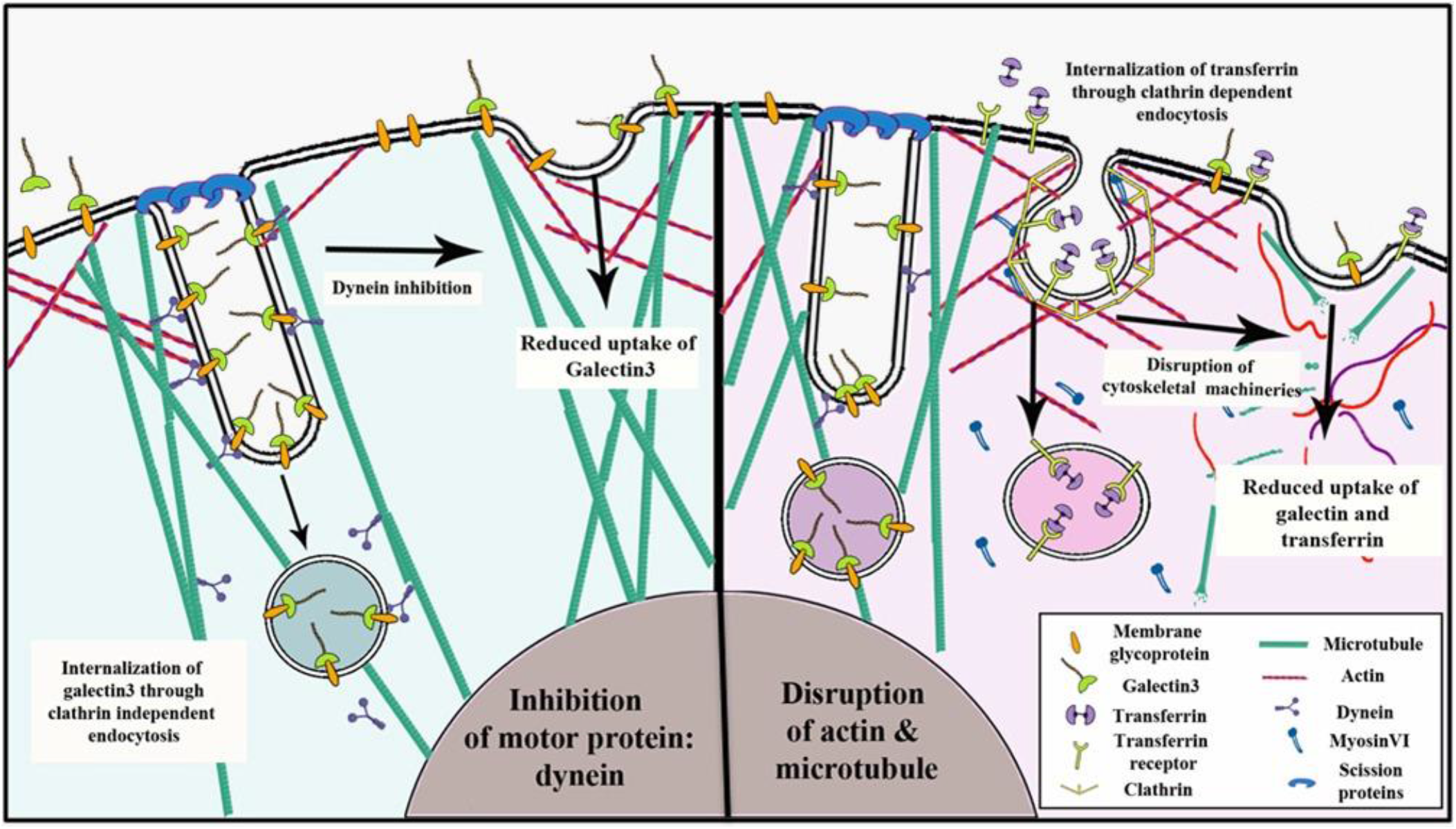
Schematic representation of the outcome of our current study: Dynein motor is recruited on tubular invaginations of Gal3 induced plasma membrane invaginations whose length decreases upon dynein inhibition. The cargoes of Gal3 such as CD98 and CD147 are also found to be affected upon dynein inhibition.

## Materials and methods

### Cell culture reagents

Dulbecco’s modified Eagle’s medium (DMEM), Hams-F12 media were purchased from Lonza. Fetal bovine serum (FBS), penicillin-streptomycin, trypsin-EDTA (0.25%), phosphate buffer saline (PBS) was purchased from Gibco. Transferrin-A561, Transferrin-A488 and Transferrin-A647, and phalloidin A488 were purchased from Invitrogen. Cell culture dishes for adherent cells (treated surface), Triton X-100 were purchased from Himedia. Petri dishes and culture plates were obtained from Tarsons. Paraformaldehyde (PFA) was purchased from Merck. DMSO was ordered from SRL. Collagen Type I was purchased from Corning. Hoechst 34580 were purchased from Thermo Fisher. FITC conjugated CTxB, insulin and dihydrocortisone were obtained from Sigma.

### Cell Culture

Mouse embryonic fibroblasts (MEFs), MDA-MB-231, Retinal Pigmented Epithelial cells (RPE1) and HeLa and were cultured in Dulbecco’s Modified Eagle medium (DMEM) with 10% fetal bovine serum, 2mM glutamine, 100U/ml penicillin-streptomycin and grown in 5% CO_2_ atmosphere at 37°C with 90% humidity. MEFs, SUM159, MDA-MB-231 and RPE1 were obtained as kind gift from the laboratory of Prof. Ludger Johannes at Curie Institut, Paris. SUM159 cells were cultured in Hams-F12 media with 5% fetal bovine serum, 2mM glutamine, 10ug/ml insulin, 10ug/ml dihydrocortisone, 100U/ml penicillin-streptomycin in 5% CO_2_ atmosphere at 37°C with 90% humidity. Cells were plated onto the coverslips for fixed cell imaging at least 24 hours prior to the experiments.

### Immunofluorescence experiments

The dilutions used for immunostaining experiments were 1:100 for primary antibodies and that of secondary antibodies were 1:1000 in case of CD98 and CD147, and 1:500 for DYNC1H1 and Hook1. Mouse monoclonal antibodies to human CD98 (clone-44D7) was from bio rad, human CD147 raised in mouse (clone-HIM6) was from Biolegend, human DYNC1H1 polyclonal raised in rabbit (Accession Q14204) was from Thermo Fisher Scientific. The secondary antibodies used were goat anti mouse IgG (H+L) Alexa Fluor 647 conjugate, goat anti mouse IgG (H+L) Alexa Fluor 488 conjugate and goat anti rabbit IgG (H+L) Alexa Fluor 488 conjugate, purchased from Thermo Fisher Scientific.

### Inhibition of Dynein

MEFs or SUM159 cells were previously seeded for 24-36 hours at 37°C, in well plates with cover slips. After reaching appropriate cell density the complete growth media was replaced with serum free media for drug treatments and uptake studies. The final concentrations of Ciliobrevin D (calbiochem) and Dynapyrazole A (Sigma) was 50uM and 15uM respectively. The Inhibitor treatments were carried out at cell culture conditions for 30 minutes and then Gal3, Gal8, CTxB or Tf were incubated in the presence of the inhibitors for an additional 15 minutes. The cells were washed with PBS or PBS++, fixation done with 4% PFA for 10 mins at 37°C and the coverslips were then mounted on a glass slide with mowiol, containing DAPI or Hoechst for confocal imaging.

### ATP depletion assays

ATP depletion was carried out by pre-incubating the cells with ATP depletion medium for 15 minutes, consisting of 10mM 2-deoxy-D-glucose and 10mM NaN3, PBS++ (PBS+0.5mM CaCl2+0.5mM MgCl2). This was carried out at 37’C. The controls were kept at serum free medium. The uptake of endocytic markers was carried out in the presence of ATP depletion media for an additional 15 minutes at cell culture conditions.

### Invasion assay using MDA-MB-231 3D spheroid model

MDA-MB-231 spheroids were formed using the hanging drop method. From the homogenous cell suspension with the concentration of 5000 cells/ml, droplets of cells were placed underneath the lid of the petri dish filled with 1X PBS for humidity. The cell droplets were incubated at 37 °C for 24 h to facilitate 3D spheroid formation. After 24 h, the spheroid formation was visualized using an optical microscope with 10X objectives. The spheroids were dispersed gently onto the collagen-DMEM complete media matrix (in 3:1 vol/vol ratio). The spheroids placed in the matrix were incubated at 37°C for 1 h. For the treatments, the spheroids were supplemented with DMEM complete media and treated with ciliobrevin D, Dynapyrazole A, transferrin, galectin-3 for 24 h at 37°C. The control set was supplemented only with DMEM complete media. Five to seven spheroids were taken for each treatment set. After 24 h, the spheroids were fixed with 4% PFA at 37°C for 20 mins. The spheroids were permeabilized with 0.1% Triton X-100 for 30 mins and stained with 0.1% phalloidin A488 for 1 hour. Followed by staining, it was washed with 1X PBS and mounted. Further, the spheroids were visualized using a confocal microscope with a 10X objective by exciting at 488nm.The invasion index of the cells from the spheroid was observed with respect to control, and the invasion index was calculated using the following equation *Invasion index=Distance migrated by the cells from the surface of the spheroid/diameter*.

### Dynein Heavy chain knockdown

Knockdown of DYNC1H1 subunit of dynein was achieved by using mission® esiRNA from Sigma (EHU059651). The esiRNA was prepared according to the manufacturer’s instructions and the cells were transfected with the esiRNA pool with Hiperfect (Qiagen). Transfection was performed according to the manufacturer’s instructions. Knockdown was performed in MEFs and the final concentration used was 60nM for the period of 48 hours. Prior to siRNA treatment, cells were incubated in cell culture conditions for 12-24 hours.

### Expression and purification of recombinant Gal3 and Gal8 and their conjugation with fluorophores

His-tagged Gal3 and His-Tagged Gal8 (pHis2Parallel) were kind gifts from the Johannes team at Institut Curie, Paris. The above two plasmids were expressed in Rossetta2-pLysS strains in LB media. Protein expression was induced at OD600nm 0.8 with 60 μM IPTG at 21°C overnight. The bacterial pellet was resuspended in ice-cold PBS containing 10mM Imidazole, protease inhibitor cocktail, 10mM lactose, 2mM TCEP and 10% Glycerol. His-tagged affinity purification was achieved using Cobalt-resin (Pierce) and gel filtration (Superdex75 10/300). For affinity purification elution was carried out using 500mM imidazole with 10mM lactose in PBS. For size exclusion chromatography the buffer used for elution was PBS with 10mM lactose (pH 7.4). For labeling with fluorophores, 2mg/ml of proteins was used and conjugation was done with amine-reactive (NHS-ester) fluorophores as per the instructions given by Invitrogen. The unlabeled fluorophores were separated using PD-10 columns (GE Healthcare).

### Endocytosis assays

For endocytosis assays, the cells were pretreated for 30 minutes with the above-mentioned inhibitors before pulsing the cells with endocytic markers. The concentrations used for Gal3, Gal8, Transferrin, CTxB used were 5μg/ml. The cells were incubated for 15 mins and then washed with PBS and ice-cold Glycine washes as per the reference (Lakshminarayan et al., 2014). Ice-cold Glycine washes (pH 2.2) were given in order to remove the cell surface markers wherever necessary. Then the cells were fixed with 4% PFA and mounted using Mowiol with DAPI (Sigma). Two independent experiments were performed to confirm the results.

### Confocal microscopy

Confocal microscopy was carried on Leica Laser Scanning Confocal microscope (TCS SP8) using 63X/1.4 NA, apochromatic oil immersion objective. DAPI and Hoescht were excited using UV laser at 405nm. Alexa-488 and FITC conjugated proteins were excited using 488nm line Argon laser. Cy3 and Alexa-546 conjugated proteins were excited using 561nm DPSS laser. For Cy5 and Alexa-647 conjugates HeNe laser at 633nm was used for excitation. The detectors used were Hybrid and PMT.

### Image analysis and Statistical analysis

Maximum intensity projection of Z stacks were used for quantification of cells and raw intensity density was calculated. To show perinuclear localization, mid-stacks were used for quantification. The images were processed using Fiji Image J 2.1.0 (NIH). The difference between the control and test groups’ mean values was analyzed by the student’s t-test or one-way ANOVA using GraphPad Prism 9.0.

## Acknowledgements

We extend our sincere thanks to Dr. Christian Wunder and Prof. Ludger Johannes from U1143/UMR3666 Institut Curie, Paris for their extremely helpful guidance and discussion all throughout the project. We sincerely thank all the members of DB lab for discussions and stimulating ideas. C.M., H.N.A and P.M. thank IIT Gandhinagar, Ministry of education, Govt of India for research fellowships. D.B. thanks the Science and Engineering Research Board, Department of Science and Technology, Government of India for a Ramanujan fellowship; and the Indian Institute of Technology Gandhinagar for initial funding support. The central instrumentation facilities at IITGN are duly acknowledged.

The authors declare no competing or financial interests

## Supplementary information

**Figure S1.**
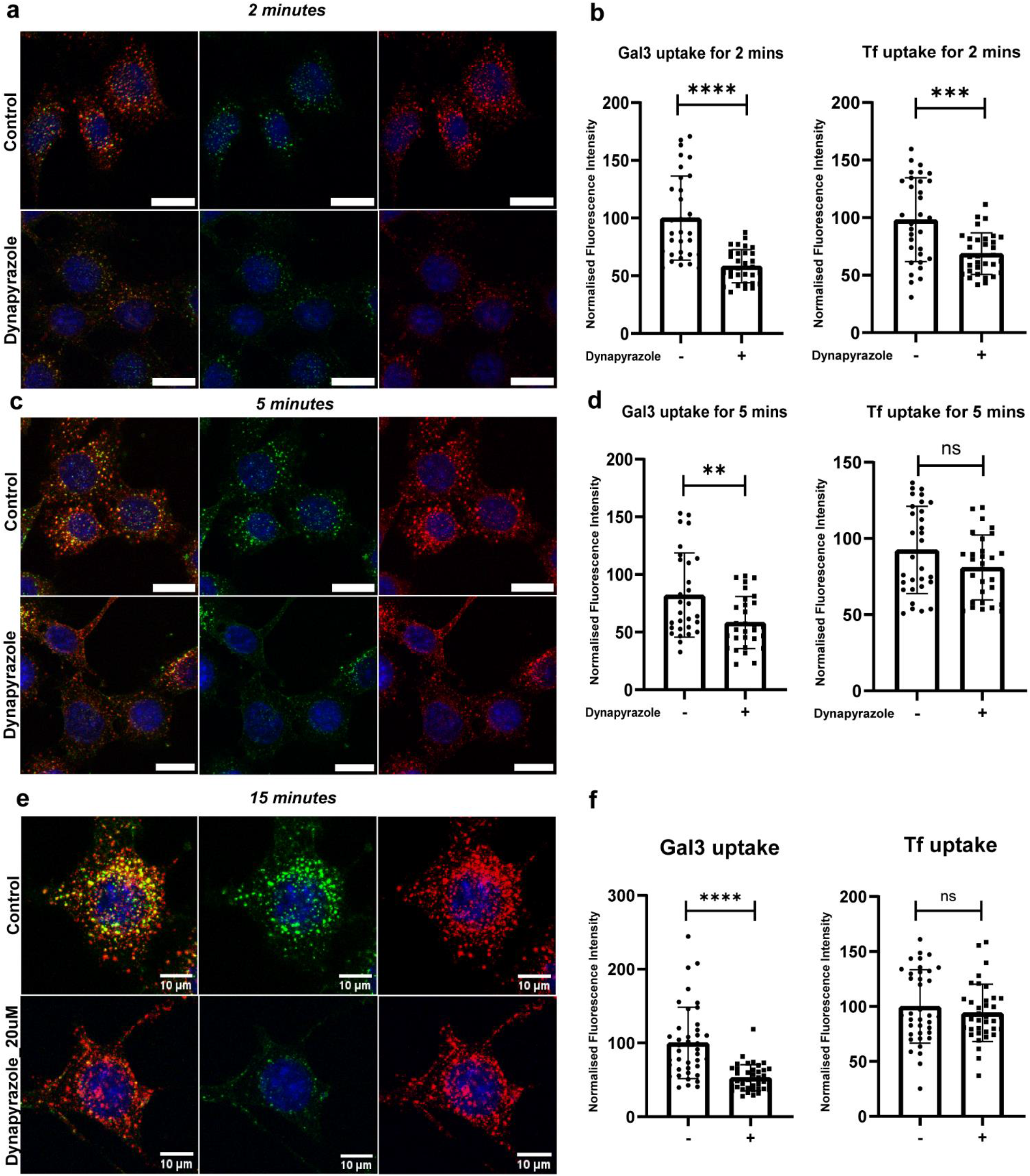
Gal3 is affected upon dynein inhibition at different time points: (a) Prior treatment of cells with Dynapyrazole followed by 2 minutes uptake of Gal3 and Tf affected the uptake of both cargoes. Dynein could be playing a role during the initial stages CME and CIE processes. (b) Quantification for Gal3 and Tf uptake for 2 minutes time interval showed significant change in Dynapyrazole treated cells vs the control. (c) The cells treated with the inhibitor affected the uptake of the cargoes at 5 minutes interval. (d) Statistical analysis of Gal3 and Tf uptake for 5 minutes time point. (e) On increasing the concentration of Dynapyrazole, we observe decrease in Gal3 uptake and Tf distribution in the cell also get affected. (f) Quantification of Gal3 and Tf fluorescence intensities shows the similar results compared to 15μM of dynapyrazole used. **±** S.D. of normalized fluorescence intensity of 30-35 cells. Asterisks denote significant difference from the control vs dynapyrazole treated, with ** p ≤ 0.01, *** p ≤ 0.001, **** p ≤ 0.0001 and ns represents non-significant difference from control.

**Figure S2.**
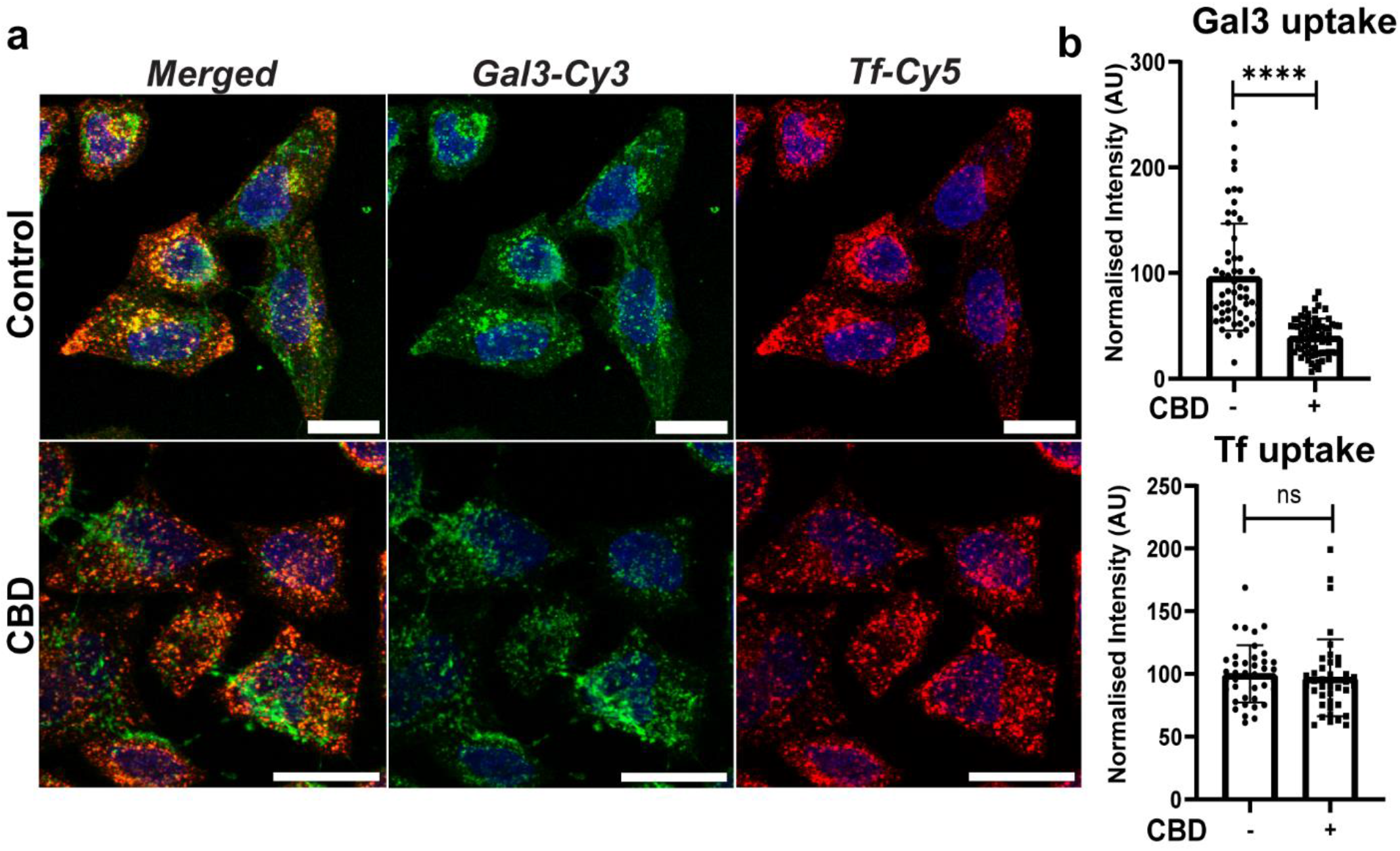
Uptake of Gal3 is affected upon CBD treatment. (a) Confocal images of MEFs showing the uptake of Gal3-Cy3 in green and Tf-Cy5 in red after 30 minutes of CBD treatment. CBD treatment reduces Gal3 uptake but not Tf. (b) Quantification of Gal3 and Tf uptake upon CBD treatment. Error bars represent standard deviation. Normalized fluorescence intensity was calculated from 35-45 cells. Student T-test was used for obtaining statistical significance with **** p ≤ 0.0001 and ns represents non-significant difference from control. Scale bar is 20μm.

**Figure S3.**
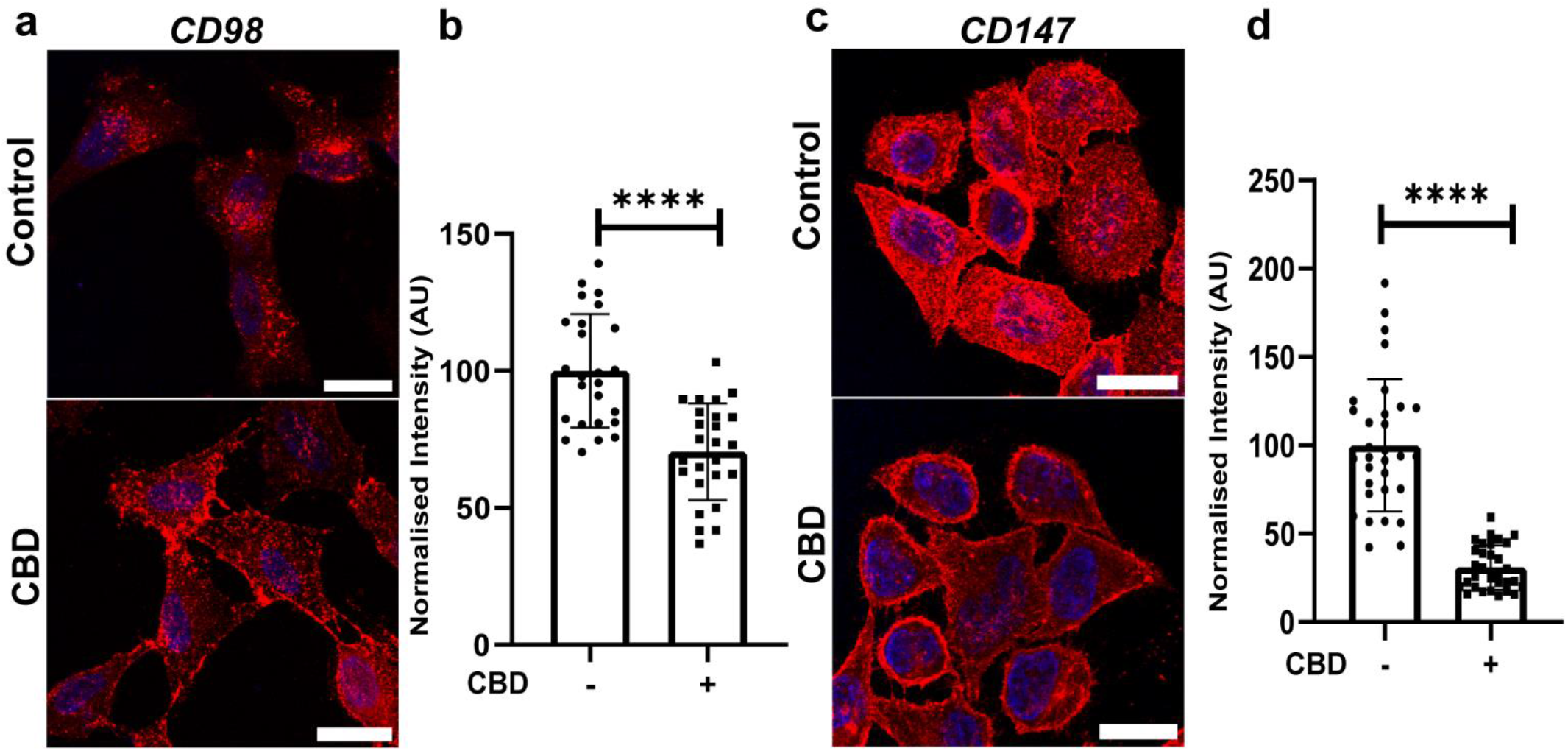
Uptake of CD98 and CD147 is affected on dynein inhibition: (a) Confocal images of SUM159 cells immunostained for CD98 using Alexafluor-647 conjugated secondary antibody (represented in red). The images show difference in the pattern of CD98 localization in cells in control vs CBD treated conditions. (b) Quantification for CD98 uptake in cells. The graph shows significant difference between control and CBD treated cells. (c) Confocal images of SUM159 cells immunostained for CD147 using Alexafluor-647 conjugated secondary antibody (represented in red). The images show difference in the pattern of CD147 localization in cells in control vs CBD treated conditions. (d) Quantification for CD147 uptake in cells. The graph shows a significant difference between control and CBD treated cells similar to CD98. Error bars represent standard deviation. Normalized fluorescence intensity was calculated from 35-40 cells. Student T-test was used for calculating statistical significance with ns being non-significant from the control. Student T-test was used for obtaining statistical significance with **** p ≤ 0.0001 and ns represents non-significant difference from control. Scale bar is 20μm.

**Figure S4.**
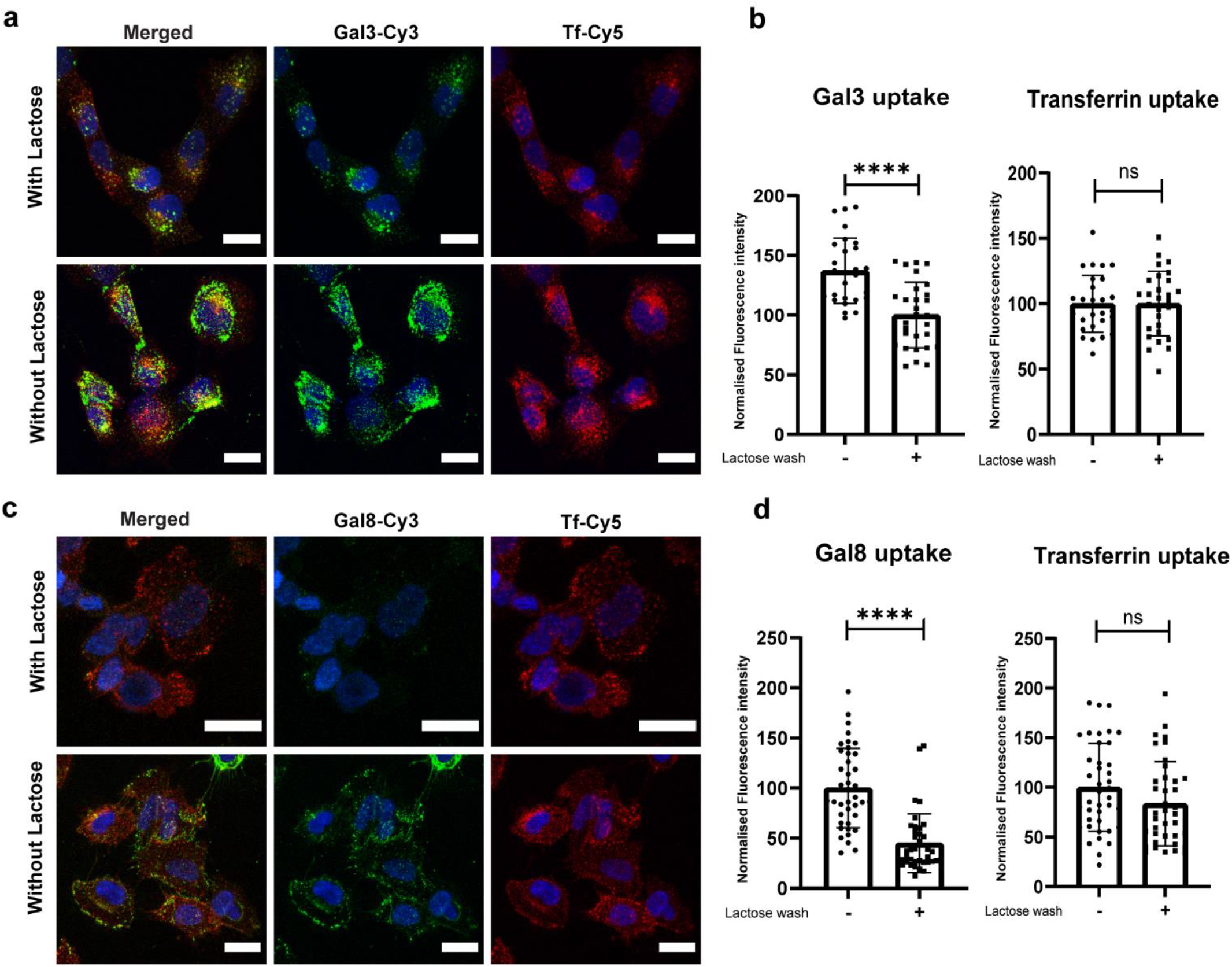
Gal3 and Gal8 are inhibited by lactose: Lactose is being used as a competitive inhibitor of galectins. Hence, we used 150 mM of lactose to study the effect of Gal3 and Gal8 uptake into the cells. (a) Pretreatment with lactose affects Gal3 binding but not transferrin. (b) Quantification of Gal and Tf uptake upon lactose treatment. (c) Gal8, another lectin was also inhibited on lactose treatment while Tf remains unaffected. (d) Quantification for Gal8 and Tf uptake upon lactose treatment. This was being used further to test the uptake of other receptors which gets internalized through CIE. S.D. of normalized fluorescence intensity of 30-35 cells. Asterisks denote significant difference from the control vs lactose treated cells, with and **** p ≤ 0.0001 and ns represents non-significant difference from control.

**Figure S5.**
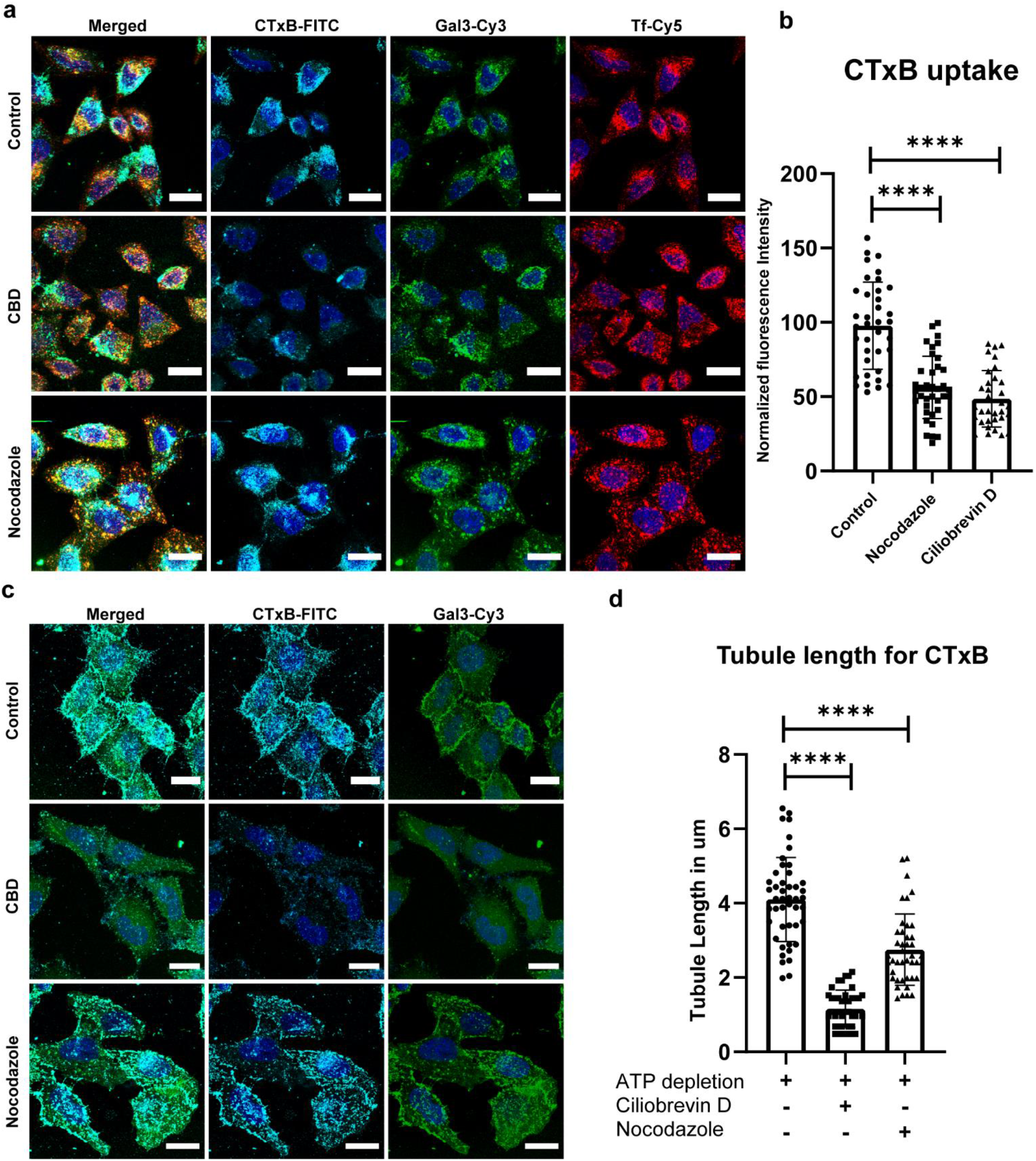
CTxB is used as model cargo for studying CIE: (a) Upon dynein inhibition and microtubule depolymerization, CTxB uptake gets affected, similar to Gal3. CBD treatment did not affect Tf uptake but nocodazole affected the uptake of Tf. (b) Quantification for CTxB uptake upon Nocodazole and CBD treatment with respect to control cells. (c) CTxB form tubules upon ATP depletion which is affected upon CBD and Nocodazole treatment. The similar pattern is observed in Gal3 as well. (d) Quantification for tubular length exhibited by CTxB for CBD and Nocodazole treated ATP depleted cells. 30 cells were considered for counting the tubule lengths.

**Figure S6.**
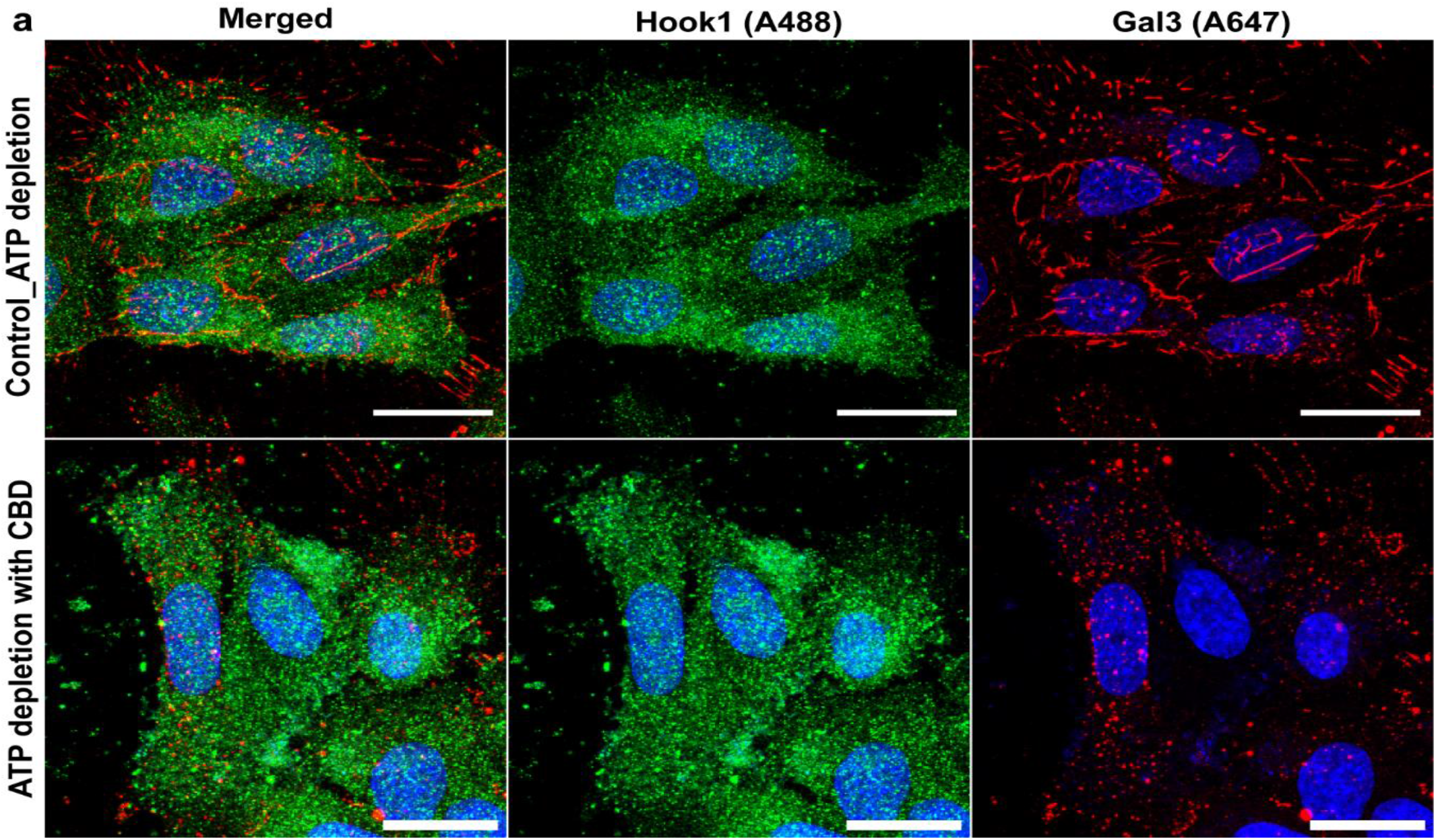
Hook1, a dynein adaptor does not co-localize with Gal3 tubules: (a) ATP depletion for 15 minutes resulted in the tabulation of Gal3 which significantly affected on dynein inhibition. However, Hook1 did not co-localize with the tubules.

## Notes

### Competing Interest Statement

The authors have declared no competing interest.

